# Deep oncopanel sequencing reveals fixation time- and within block position-dependent quality degradation in FFPE processed samples

**DOI:** 10.1101/2021.04.06.438687

**Authors:** SEQC2 Oncopanel Sequencing Working Group, Yifan Zhang, Thomas M. Blomquist, Rebecca Kusko, Daniel Stetson, Zhihong Zhang, Lihui Yin, Robert Sebra, Binsheng Gong, Jennifer S. LoCoco, Vinay K. Mittal, Natalia Novoradovskaya, Ji-Youn Yeo, Nicole Dominiak, Jennifer Hipp, Amelia Raymond, Fujun Qiu, Hanane Arib, Melissa L. Smith, Jay E. Brock, Daniel H. Farkas, Daniel J. Craig, Erin L. Crawford, Dan Li, Tom Morrison, Nikola Tom, Wenzhong Xiao, Mary Yang, Christopher E. Mason, Todd A. Richmond, Wendell Jones, Donald J. Johann, Leming Shi, Weida Tong, James C. Willey, Joshua Xu

**Author notes:** These authors contributed equally to this work (Yifan Zhang, Thomas M. Blomquist, Rebecca Kusko). Corresponding authors: Dr. James C. Willey, Dr. Joshua Xu.

## Abstract

Clinical laboratories routinely use formalin-fixed paraffin-embedded (FFPE) tissue or cell block cytology samples in oncology panel sequencing to identify mutations that can predict patient response to targeted therapy. To understand the technical error due to FFPE processing, a robustly characterized normal cell line was used to create FFPE samples with four different pre-tissue processing formalin fixation times. A total of 96 FFPE sections were then distributed to different laboratories for targeted sequencing analysis by four oncopanels, and variants resulting from technical error were identified. Tissue sections that failed more frequently showed low cellularity, lower than recommended library preparation DNA input, or target sequencing depth. Importantly, sections from block surfaces were more likely to show FFPE-specific errors, akin to “edge effects” seen in histology, and the depth of formalin damage was related to fixation time. To assure reliable results, we recommend avoiding the block surface portion and restricting mutation detection to genomic regions of high confidence.

## Introduction

Next generation sequencing (NGS) is now an integral tool in the “precision” cancer care arsenal (NCI-match studies; ECOG and other substudy trials). Despite excellent performance for somatic mutation calling^1^, preanalytical variation continues to limit the quality and quantity of cancer specimens, and this ultimately impacts NGS accuracy and reproducibility^2–4^. One source of preanalytical error stems from formalin-fixation and paraffin-embedding (FFPE). FFPE processing of tumor specimens is central to histologic diagnosis of cancer and its pursuant sub-classification, grading, staging, and adequacy-assessment for ancillary studies in routine clinical workup^5^. For this reason, many ancillary prognostic and treatment markers are optimized for FFPE tissue^6,7^. When combined with a growing trend of limited specimen size and quality^8^, in many circumstances, tissue is processed for FFPE to meet the bulk of current standard care needs^5^. However, FFPE processing harbors substantial and highly variable effects on nucleic acid quality and quantity^9^. Thus, platforms for targeted NGS analysis of FFPE clinical specimens must be subjected to rigorous analytical validation^10–12^. It is of utmost importance that we understand factors that influence the reproducibility and accuracy of NGS testing in FFPE tissue to ensure that cancer care is fully optimized.

The process of formalin fixation and paraffin embedding includes several steps with varying degrees of: 1) fixation, 2) progressive dehydration, 3) clearing, and 4) molten paraffin infiltration^13^. For each step, clinical labs independently establish the ideal amount of time, temperature, pressure, agitation, and reagent composition, typically with the primary goal of optimizing histologic quality^13,14^. Some guidelines do exist for specific preanalytical FFPE metrics (e.g., fixation time). For example, the College of American Pathologists recognizes that fixation time can influence accuracy and reproducibility of ancillary tests performed on FFPE such as ER/PgR immunohistochemistry and HER2 in-situ hybridization; and for these specimens, formalin fixation should be limited to >6 or <72 hours of total formalin exposure^6,7^. Still, false somatic mutation calling rates by NGS vary greatly for FFPE specimens from different, and even within the same, anatomic laboratories^15^. Of even greater concern, FFPE-derived sequencing errors may arise at clinically relevant loci and at actionable allelic frequencies^16^. Working groups are beginning to formulate and extend more detailed recommendations on pre-analytical controls^4^. While it is important that these practice recommendations are based on human biospecimen research data, some important questions may benefit from additional studies using carefully contrived sample sets as conducted here.

In this study, the Oncopanel Sequencing Working Group of the FDA-led Sequencing Quality Control Phase II (SEQC2) consortium extended the study^1^ to investigate the impact of FFPE processing. This line of inquiry has served to address the insufficiency of well-controlled data regarding FFPE tissue, its preanalytical metrics, and impact on NGS accuracy and reproducibility^17^. Here, we adopted an easy to follow FFPE preparation protocol (see Methods for details) that is highly analogous to clinically obtained cell block cytology sample processing and can be readily applied to existing reference cell line materials^18–20^. The benefits of the approach used here are that: 1) reference variant data and the ability to compare with numerous orthologous molecular methods provide robust information for accuracy and reproducibility studies, 2) genetic heterogeneity across replicate measures is minimized (a common challenge in clinical FFPE NGS studies), and perhaps most importantly, 3) preanalytical FFPE effects on technical artifacts can be well-documented and controlled, enabling future meta-analysis and subsequent guideline recommendations. As an initial investigation, cultured cell samples from a normal cell line^18^ (Agilent Male Lymphoblast Cell Line) were subjected to varying formalin fixation times between 1 and 24 hours prior to tissue processing^4,21,22^. FFPE sectioning samples at multiple locations within each FFPE block were selected and distributed to four independent laboratories for targeted NGS following amplicon or hybrid capture enrichment. Based on reference variant data, we identified false positive (FP) calls and estimated FP rate (FPR) within each panel’s targeted region—a key quality control metric demonstrated first in our cross oncopanel investigation^1^. Comprehensive analysis was then conducted on FP calls and FPR to decipher the effects of FFPE factors including formalin fixation time and tissue block section position.

## Results

### Overview of study design and analysis

To investigate the effect of formalin fixation time and tissue block section position on targeted NGS analysis of FFPE specimens, we designed a comprehensive study querying several key components. **Fig. 1** displays the flow of three major components: FFPE sample preparation (**Fig. 1A**), sequencing experiments with four diverse oncopanels (**Fig. 1B**), and data quality control and analysis (**Fig. 1C**). High quality genomic DNA from a single normal cell line (Agilent Male Lymphoblast Cell Line) was sequenced with multiple oncopanels and technical replicates in our companion study^1^. These datasets enabled the establishment of a known variant set for each panel and the subsequent detection of artifacts induced by the FFPE process (**Fig. 1C**). Cultured cell populations were used to make FFPE samples with four different formalin fixation times. Equal amounts of cultured cells were mixed in each gel matrix mold, followed by formalin fixation, routine tissue processing, and paraffin embedding (**Fig. 1A**). Samples were created by sectioning FFPE blocks as described in the Methods section. Based on their estimated cell counts and positions in the FFPE blocks, samples were assigned to two categories: surface (either top or bottom) or inner FFPE samples (see Methods for details). Each laboratory extracted and quantified genomic DNA from 24 samples evenly distributed across eight distinct FFPE blocks. NGS sequencing experiments and subsequent bioinformatics processing were conducted following vendor recommended protocol (**Fig. 1B**, see Methods). These FFPE samples were sequenced by four oncopanels (**Table S1**): AstraZeneca 650 genes Oncology Research Panel (AZ650), Burning Rock DX OncoScreen Plus (BRP), Illumina TruSight Tumor 170 (ILM), and Thermo Fisher Oncomine Comprehensive Assay v3 (TFS). Variant calling results and QC data were collected and submitted to the Working Group for integrated analysis (**Fig. 1C**, see Methods for details).

**Figure 1.**
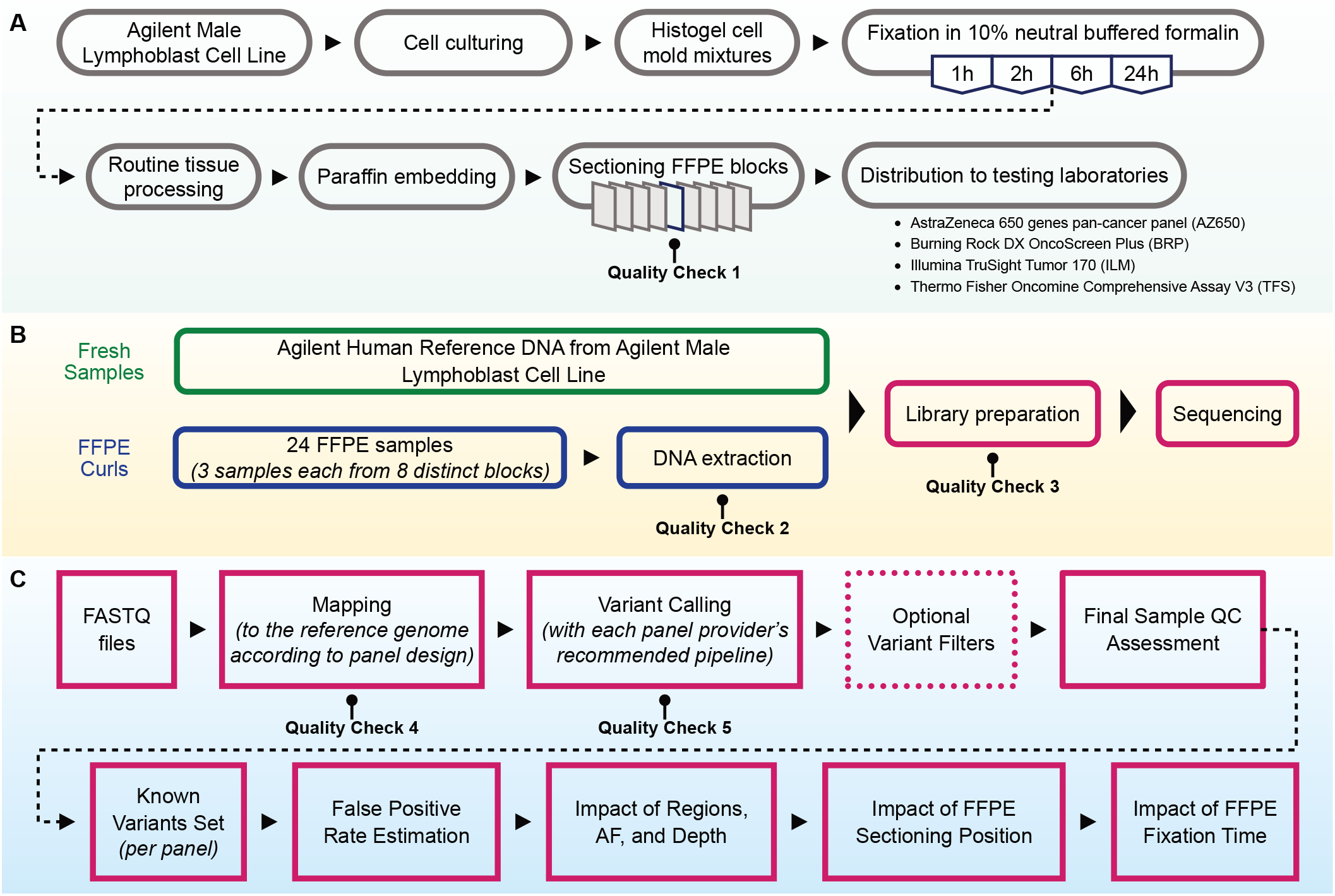
Overview of study design. (**A**) FFPE sample preparation workflow with four different formalin fixation times: 1-, 2-, 6-, and 24 hours. (**B**) Oncopanel sequencing experiments with in-laboratory DNA extraction. (**C**) Panel-specific variant calling followed by uniform and integrated analysis to assess the impacts of genomic region, VAF threshold, depth, formalin fixation time, and sample position in the FFPE block.

Five QC check steps were included in our workflow (**Fig. 1**): 1) cell count estimation, 2) DNA extraction yield, 3) library construction yield, 4) median depth and library complexity computed from mapped reads, and 5) variant histograms by variant type and VAF for the detection of oxidative damage or contamination. All QC data are provided in **Supplementary File 1**. Six samples failed in the ILM experiments based on very low target sequencing depth and low library yield, which may be related to the lower than recommended library input. Four samples failed both attempts of library preparation for TFS and were not sequenced. In each case, those failed samples appeared to have been processed together in one batch, potentially failing at the same step between DNA extraction and library construction. Five samples were possibly contaminated before or during the NGS experiments (two in AZ650, two in BRP, and one in ILM, respectively). These samples were excluded from further analysis on FFPE effects.

### The main constituents of the known variant set were homozygous or heterozygous germline variants

To differentiate FPs from true variants, it was imperative to first build the set of reference (known) variants for each panel. By aggregating the variants called by over 75% of the fresh gDNA samples that were free of FFPE damage, we generated a set of known variants for each panel. The lone exception was AZ650 whose known variant set was generated from FFPE samples that passed stringent QC filters because AZ650 was not included in our companion study of multiple oncopanels using fresh gDNA samples^1^. To determine FPs introduced by FFPE processing, variant calls from each FFPE sample were compared with the known variant set for each respective panel.

We expected to detect only homozygous and heterozygous variants in the known variant sets since a normal cell line was used. In general, the variant allele frequencies (VAFs) were close to either 0.5 or 1, with heterozygous variants (near 0.5) showing more dispersion in allele frequency than homozygous variants (**Fig. 2A**). Variants in the known variant sets located outside of the consensus high confidence targeted region (CTR, see Methods for details) showed greater VAF dispersion than those within the CTR (**Fig. 2A**), particularly for the BRP, ILM, and TFS panels. As expected, more reference variants of low VAF were reported for AZ650 than for the other three panels.

**Figure 2.**
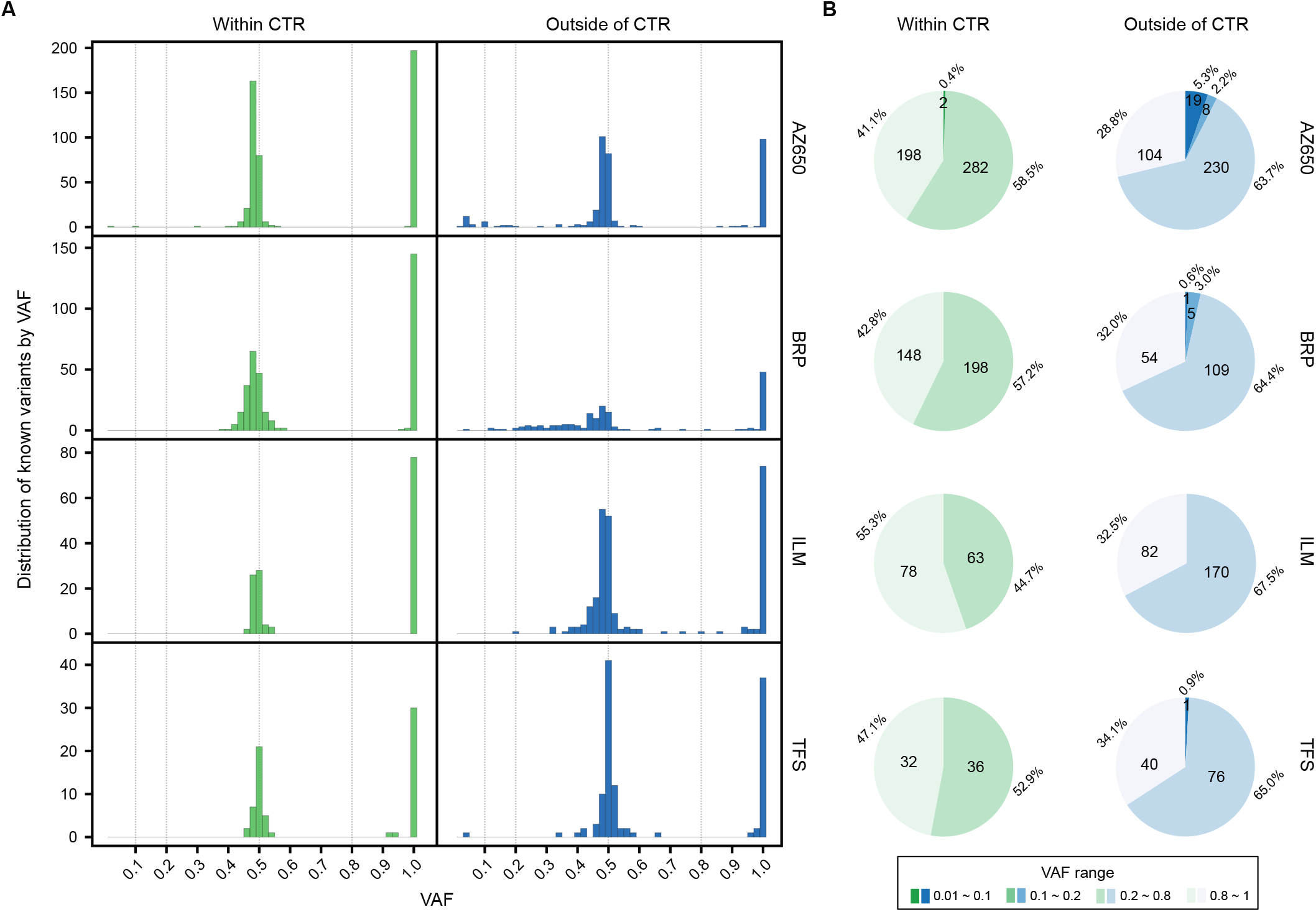
Histogram and pie chart distributions of known variants across VAF ranges confirming that most were homozygous or heterozygous germline variants. (**A**) Distribution of the number of known variants within (green) and outside (blue) of the consensus high confidence targeted region (CTR). (**B**) Number and percentage of known variants across four VAF ranges.

The distribution plot revealed few variants falling between VAF 0.1-0.3 and 0.7-0.9 (**Fig. 2A**). Thus, we choose 0.2 and 0.8 as the boundaries for heterozygous and homozygous variants. We classified the known variant set into four groups by the VAF value: homozygous variants (VAF > 0.8), heterozygous variants (0.2 < VAF ≤ 0.8), and two additional low VAF ranges separated by a VAF at 0.1. For variants called within the CTR, only two variants were in the two low VAF ranges for AZ650 while none were called for the other three panels. Among the variants called outside of the CTR, slightly more fell into the low VAF ranges and their proportion relative to all variants remained low (**Fig. 2B**). Furthermore, we confirmed the high concordance (>98%) among known variants across panels, thus the known variants were reliable. The known variants for each panel are listed in **Supplementary File 2**.

### Multiple factors can lead to a high false positive rate in FFPE samples

Oncopanel experiments can be marked as failed or flagged for further examination by collecting many quality measurements using pre-analytical and analytical instruments, bioinformatic QC tools, and reference thresholds from large cohorts. In our analysis, several QC measurements (**Fig. 3**) were found practically useful to determine whether the experiments failed or to discover outliers that required further investigation. These QC measurements, including cell count, DNA input, deduplicated sequencing depth, library complexity, G:C to T:A transversion count, and VAF distribution, are described below in detail. Cell count was an essential measurement for the quality of sequential FFPE sample sections and was the very first measurement for our pre-analytical QC check. Samples with a lower cell count usually showed a much higher FPR (**Fig. 3A**, circled in a red dashed line). However, several samples were found to have sufficient cell counts but high FPR (**Fig. 3A**, circled in an orange dashed line). In subsequent QC evaluation, these experiments were identified as failed or were flagged as outliers. Next, DNA input was another important measurement, particularly useful prior to library preparation. For example, four TFS libraries with DNA input lower than 2.5 ng showed an FPR (per million bases) greater than 50 (**Fig. 3B**). As seen in **Fig. 3B**, slightly lower DNA input substantially increased FPR. Mapping of FASTQ files to the reference genome enabled calculation of additional QC measurements. Lower sequencing depth usually correlated with a higher FPR, especially in the regions outside of the CTR (**Fig. 3C**, showcased by ILM data but observed across all panels). In addition, library complexity was a similar QC measurement for assessing the extent of PCR-assodated read duplication. Taking BRP results as an example, the FPR significantly increased when the library complexity was lower than 0.25 (**Fig. 3D**). After variant calling, several measurements can be calculated and visualized for discovering outliers. For example, G:C to T:A transversion count is a known indicator of oxidative nucleotide damage^23^. Higher G:C to T:A transversion count may be the natural state of a sample, or it may be due to exposure to oxidative radicals. Since all samples were created from a normal cell line, the samples with a high G:C to T:A transversion count were flagged as failed (**Fig. 3E**). As we were plotting the VAF histogram for each sample, we surprisingly found a handful experiments with an unexpected tail on the left side of 100%. Homozygous germline mutations are expected to be observed with a VAF of 100%, with very little or no leftward shift. Taking sample 6_G_7 as an example, the wider shift may have been the result of a small amount of contamination (**Fig. 3F**). In summary, we demonstrated how multiple QC checks can be utilized to understand the factors contributing to high FPR for some FFPE samples.

**Figure 3.**
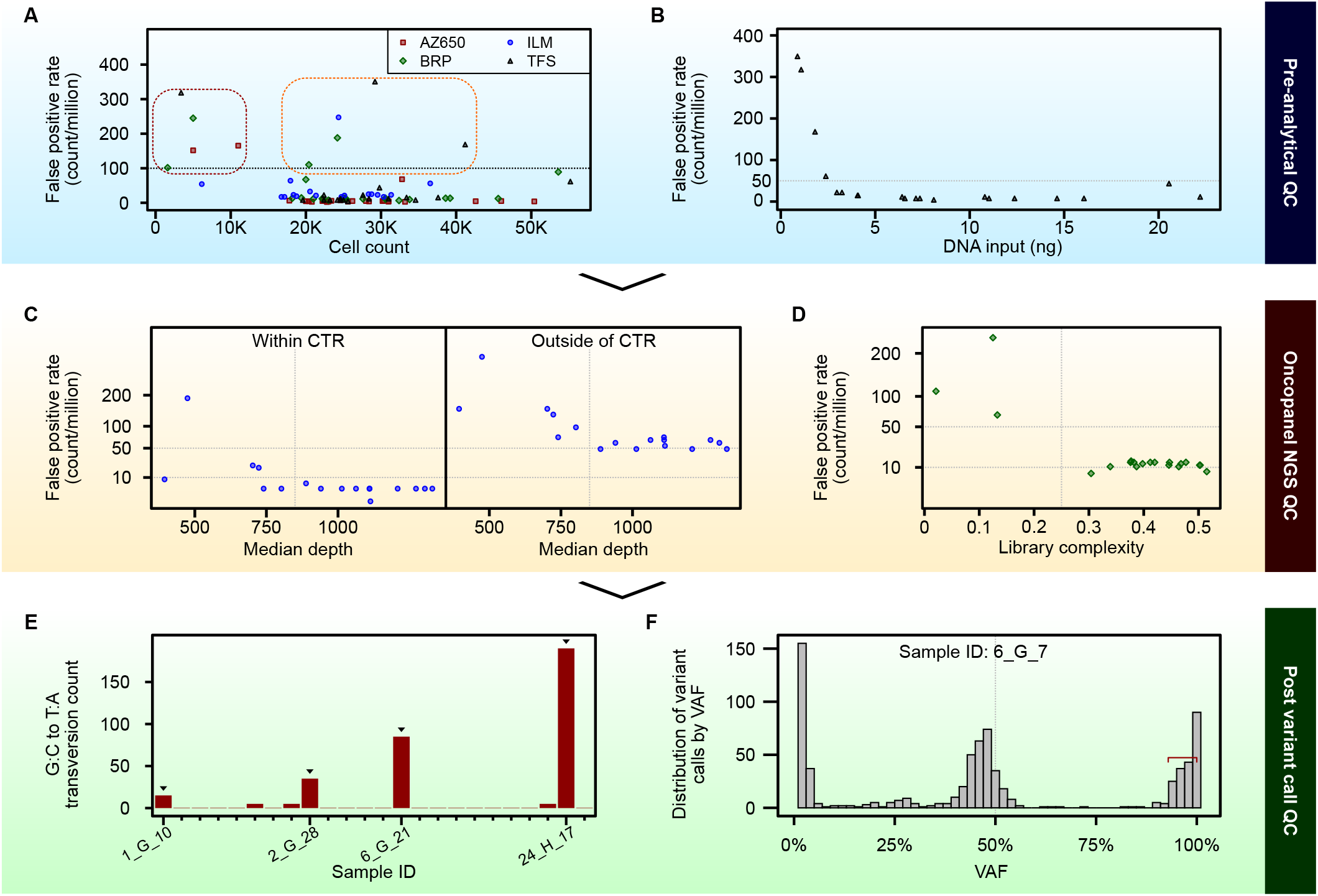
Illustration of complementary QC checks for identifying bad experiments and outliers. QC checks can be done during three major processes: pre-analytical, oncopanel NGS, and post variant calling. Multiple measurements can be used to evaluate the quality of the experiments and the data. Here we showcase a few measurements which are important in our study. (**A**) Experiments (circled with a red dashed line) with lower cell count (x-axis) usually showed a higher false positive rate (FPR, y-axis). Some experiments (circled with an orange dashed line) with cell counts (>20K) and a higher FPR failed one or more subsequent QC checks. (**B**) Experiments with low DNA input (x-axis) show high FPR (y-axis). (**C**) Lower sequencing depth (which can be measured by median depth, total depth) tends to result in higher FPR. Regions out of CTR are more affected. (**D**) Experiments were considered to have failed if library complexity was too low (below 0.25 as observed in our data). (**E**) G:C to T:A transversion count could be used to identify outliers for further investigation. (**F**) In this VAF histogram, a long tail from the left side of 100% VAF was observed in sample 6_G_7, which indicates potential contamination.

### False positive rates were impacted by genomic regions and can be effectively controlled by a VAF cutoff

The majority of surface FFPE samples failed one or more of the QC checks described above. Understandably, the surface FFPE samples were exposed to longer and more intensive chemical damage than the inner FFPE samples. Using the inner FFPE samples that passed all QC checks, we evaluated their FPRs compared against fresh samples to investigate the impacts of genomic regions and VAF cutoffs on the FPR. Given that variant detection performance may be influenced by the chosen alternate allele depth (ADP) threshold for variant calling, we also evaluated the combination of VAF cutoffs with additional ADP cutoffs. All panels showed the lowest FP rates within the CTR compared against outside the CTR (**Fig. 4A**).

**Figure 4.**
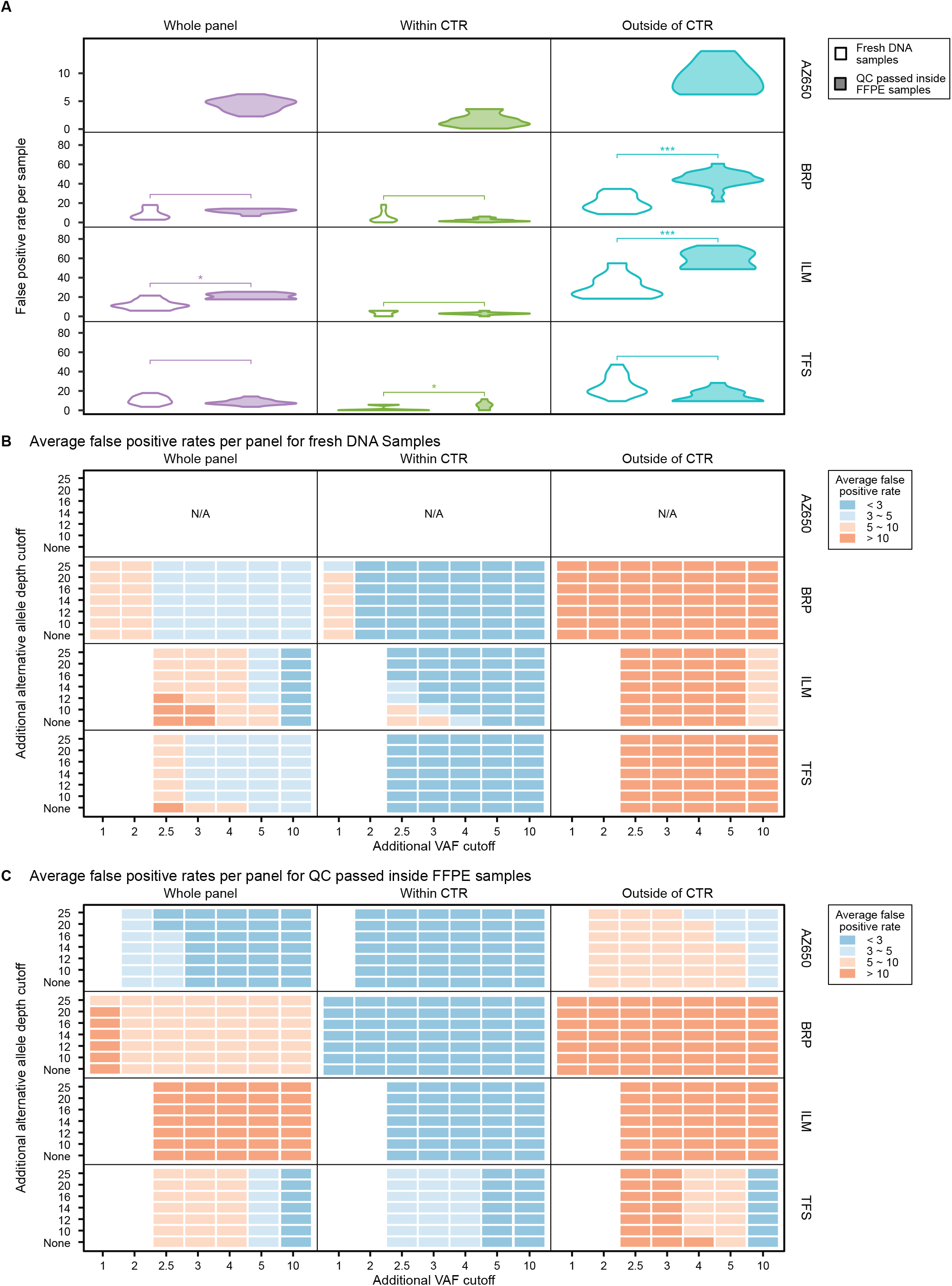
Similar false positive rates (per million base) were achieved by QC-passed inner FFPE samples in comparison to fresh DNA samples. (**A**) Violin plots of the false positive rate for fresh DNA samples versus QC-passed inner FFPE samples in three panel regions (whole panel, within the CTR, or outside of the CTR), asterisk symbols represent the significance level of the true difference in means is less than 0 (*: p< 0.05, **: p< 0.01, ***: p< 0.001). (**B**) Average false positive rate in different panel regions for fresh DNA samples after applying various additional VAF and alternative allele depth cutoffs. (**C**) Average false positive rate in different panel regions for QC-passed FFPE samples after applying various additional VAF and alternative allele depth cutoffs.

We then employed a one-tailed t-test to check whether the FPR of the QC-passing inner FFPE sample group was higher than that of the fresh sample group (**Fig. 4A**). For the whole panel region, ILM was the only panel with higher FPRs in FFPE samples. That may be explained by the lower DNA input amount (≤ 30 ng) for the FFPE samples in comparison to 50 ng for fresh DNA samples. Within the CTR, only the TFS panel showed a significant difference in FPR between the two sample groups (**Fig. 4A**). Again, the variable DNA input amounts may have been a contributing factor. Taken together, these observations suggested no consistent impact on the FP rate by the FFPE process. The standardized FFPE procedure together with the rigorous quality control checks described above could thus achieve an FP rate close to that of fresh gDNA samples.

We also applied a range of additional VAF and ADP cutoffs beyond each panel’s internal thresholds to discern the influence of VAF and ADP thresholds on FP rates (**Fig. 4B-C**). Across all of the panels, applying both criteria reduced FP rates as expected. VAF cutoffs ranging from 1% to 10% led to a higher degree of FP rate reduction than applying additional ADP cutoffs. Of note, the benefits were not consistent across sample types (FFPE samples vs fresh samples) or oncopanels. When considering FFPE samples only (**Fig. 4C**), an increase in VAF cutoff from 2.5% to 10% resulted in only modest changes in BRP’s FPR but notable changes in TFS’s FPR. The VAF and ADP cutoffs were interdependent criteria; how much each one influenced reduction of the FPR depended on analytical filters such as total sequencing depth. Based on the results of this study, we suggest that cutoff selection should be based on the desired or required FPR.

These results suggested that the CTR is a more reliable region for variant reporting, as evidenced by the much lower FPRs achieved in the CTR. Additionally, we confirmed the effectiveness of stricter VAF cutoffs to control the FFPE sample FP rate.

### Surface FFPE samples showed significantly more FFPE damage and artifacts due to hydrolytic deamination

We extended the analysis of FP variant calls to include “surface” FPFE samples and QC-failed “inner” samples. Since the FPR was much higher outside the CTR, the analysis was confined to the CTR. The FPs were further classified into four variant categories: 1) indels, 2) hydrolytic deamination introduced artifacts (G:C>A:T SNVs), 3) oxidative damage artifacts (G:C>T:A SNVs), and 4) other FP calls. The FP count per sample was plotted for each sample category (**Fig. 5A**).

**Figure 5.**
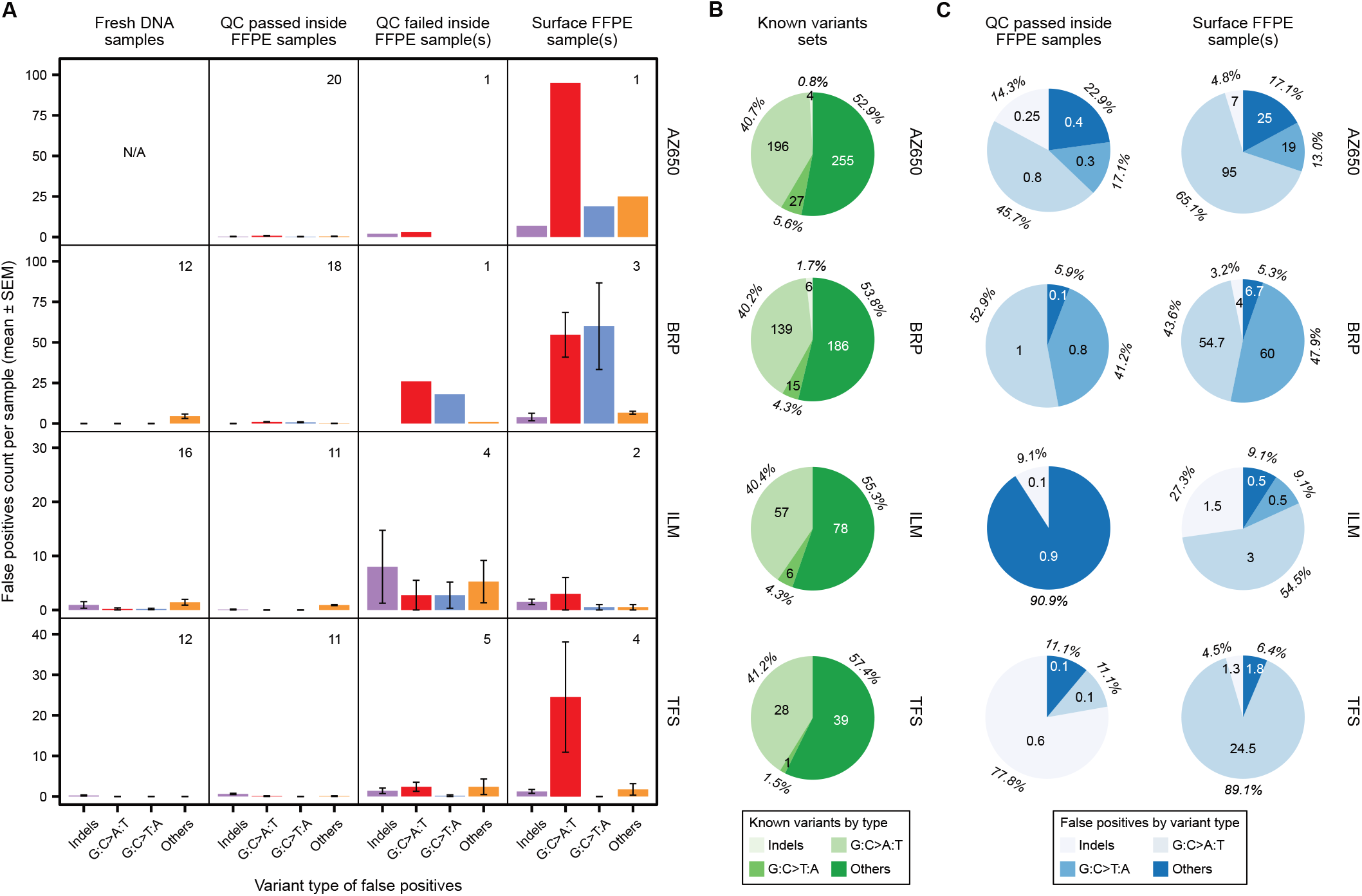
Counts and distribution of false positive calls by variant types within the consensus targeted region (CTR) indicated more FFPE damage in surface FFPE samples. (**A**) The average number of false positive calls is plotted with standard error of the mean (SEM) for four variant types within the CTR for the fresh DNA and various FFPE sample groups. The variant types were 1) indels, 2) hydrolytic deamination introduced artifacts (G:C>A:T transitions), 3) oxidative damage artifacts (G:C>T:A transversions), and 4) other FP calls. The number of samples for each sample group is inserted at the top right corner of each subplot. (**B**) Number and percentage of known variants within the CTR are shown in a pie chart over four variant types for each panel. (**C**) Average number and percentage of false positive calls per sample within the CTR are shown in a pie chart over four variant types for each panel and two sample groups (QC-passed inner FFPE samples vs surface FFPE samples).

Overall, compared with fresh and QC-passed inner FFPE samples, both surface and QC-failed inner FFPE samples produced more FP calls. Except for ILM, the surface FFPE samples consistently made more FP calls per sample than the QC-failed inner FFPE samples (**Fig. 5A**). Furthermore, across all panels, the surface FFPE samples yielded more hydrolytic deamination artifacts per sample than the QC-failed inner FFPE samples. This clear pattern indicated that the main driver of surface FFPE samples FP calls was hydrolytic deamination introduced artifacts (G:C>A:T). Within the known variant set, the G:C>A:T variants constituted approximately 40% of all variants for each panel (**Fig. 5B**). In comparison, the proportion of G:C>A:T FP calls for surface FFPE samples was much higher than 40%, which was the baseline value for the known variant sets. The lone exception was BRP whose surface FFPE samples also produced a high number of FP G:C>T:A calls per sample, presumably due to some oxidative damage (**Fig. 5C**). Understandably, the surface FFPE samples were physically exposed to more intensive formalin treatment, leading to more FFPE process-induced uracil lesions^24^. Our results clearly demonstrated that more DNA damage occurred on the surface portions of the sample during the FFPE process.

It was understandable that all five surface samples with low cell counts (1600 - 6200) failed the sequencing experiments. Even after excluding them, the QC-passing rate among the surface sample group (1 out of 5) was drastically lower than that of the inner FFPE sample group (60 out of 71). The samples taken from the inner portion of a FFPE block performed significantly better than samples from the surface portion (p<0.004, Pearson’s Chi-squared test with Yates’ continuity correction).

### Longer formalin fixation reduced the portion of high-quality samples

A previous study reported that formalin fixation times longer than 24 hours could lead to more deamination artifacts^25^. In this study, we limited the fixation time to 24 hours maximum. Within the QC-passed inner FFPE samples, we examined whether the fixation time would lead to any consistent differences in FPR. This analysis was first performed for each panel (**Fig. 6A-D**). Except for TFS, there was no observable effect of formalin fixation time on the FPR of inner FFPE samples. For TFS, the longer fixation group (i.e., 6 hours and 24 hours) appeared to show more elevated FPRs than the shorter fixation group (i.e., 1 hour and 2 hours). However, when pooling all samples together for analysis, there were no significant differences in FPR between the long and short fixation groups (**Fig. S1**). Using the FPR as a quality measure for the sample, this result indicated that longer formalin fixation time had no detrimental effect on the quality of inner FFPE samples. We then calculated the average cell count of inner FFPE samples for each block and tested whether there was any difference across the fixation time groups. The FFPE blocks were made in the same dimension with similar cell densities. As expected, the average cell count showed no effect by formalin fixation time (**Fig. 6E**). Taken together, regardless of the fixation time, inner FFPE samples showed no FFPE damage and could achieve the same low false positive rates as fresh DNA samples (**Fig. 4A, Fig. 5A**).

**Figure 6.**
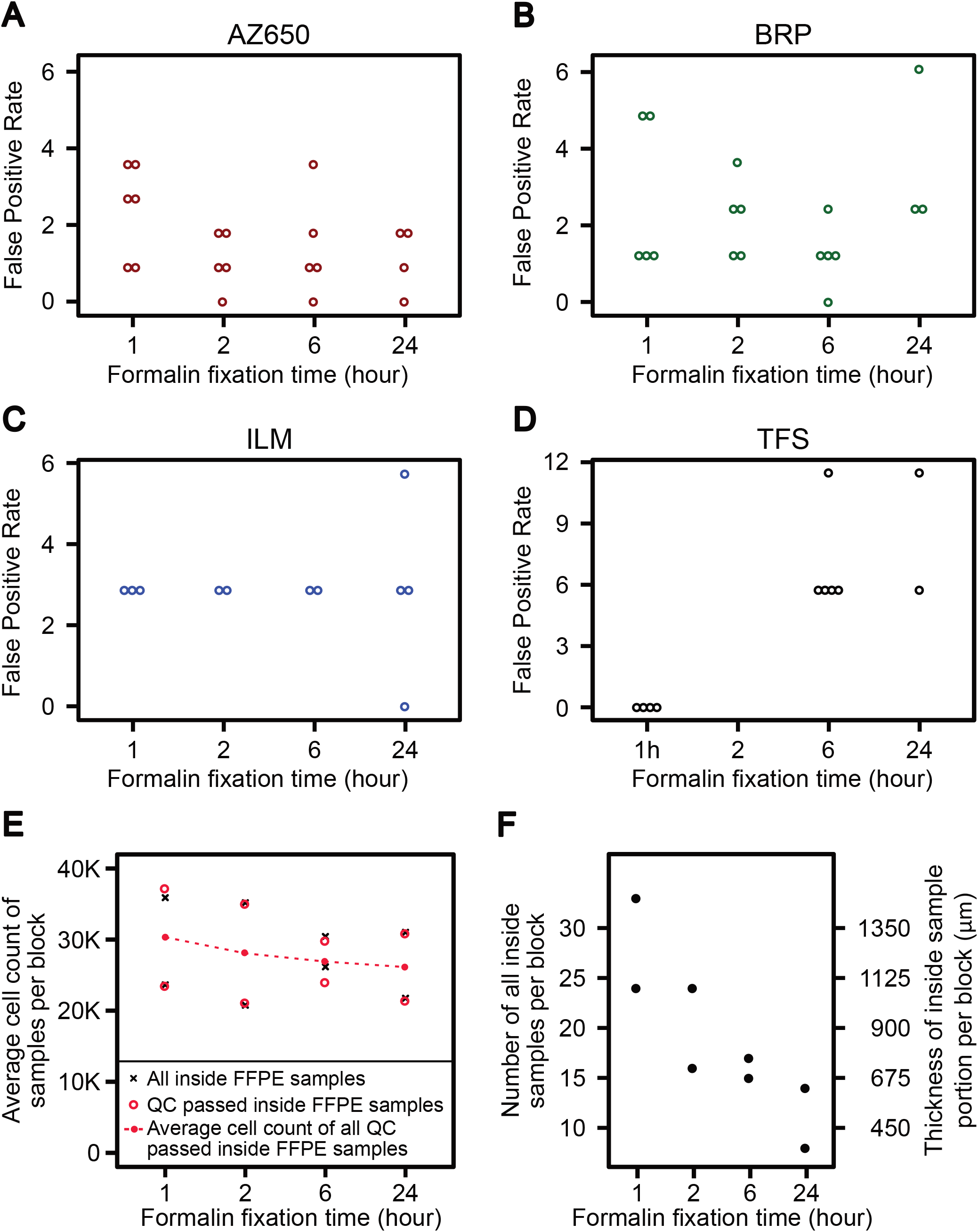
Longer formalin fixation time had no effect on the quality of inner FFPE samples but reduced their quantity. (**A-D**) For each panel, QC-passed inner FFPE samples are plotted by formalin fixation time (x-axis). Each circle represents a sample with its false positive rate per million bp shown on the y-axis. Samples are color coded per panel: red for AZ650, green for BRP, blue for ILM, and black for TFS. Except for TFS, there was no observable effect of formalin fixation time on the quality (measured by false positive rate) of inner FFPE samples. (**E**) The average cell count of inner FFPE samples remained consistent across different formalin fixation times. Two FFPE blocks were prepared under each formalin fixation time. Red circles denote the average cell counts of QC-passed inner FFPE samples taken from each block while block crosses denote the average cell counts of all inner FFPE samples within each block. These two values are shown to be very close to each other for each block. Finally, the red dots and dashed line show that the overall average cell counts are consistent across four formalin fixation times. (**F**) The quantity of inner FFPE samples is plotted for each FFPE block. The number of inner samples in each block is shown on the left y-axis while the aggregated thickness of all inner samples (i.e., thickness of inner sample portion) is shown on the right y-axis. Longer formalin fixation time clearly reduced the quantity of inner FFPE samples.

Finally, we counted all inner samples within each FFPE block and calculated their aggregated thickness (i.e., depth of the inner sample portion) within each block and compared it across the fixation time groups. Factoring the alternative sectioning slide stained for cell counting, each sample comprised nine 5 μm sections and thus was equivalent to a portion of 45 μm thickness within the block. The shorter fixation groups (i.e., 1 hour and 2 hours) clearly contained more inner samples and the inner portion was thus thicker than that in the longer fixation group (i.e., 6 hours and 24 hours) (**Fig. 6F and Supplementary File 1 – Sample Cell Counts Table**). This formalin fixation-related shrinkage artifact is well documented^5^, and this condition combined with longer fixation times could have led to fewer high quality inner FFPE samples available for oncopanel testing.

## Discussion

The goal of this study was to identify and characterize poorly understood sources of technical variation associated with targeted NGS analysis of variants in FFPE tissue and cell block samples. Through the effort of the Oncopanel Sequencing Working Group of the SEQC2 consortium, we used multiple oncopanels to systematically survey the effect of formalin fixation time, genomic location, and sectioning position within an FFPE block. Owing to this investigation’s focus on technical variation, a single robustly characterized normal cell line (Agilent Male Lymphoblast Cell Line) was used to create FFPE cell blocks with four different formalin fixation times. We adopted an FFPE procedure that is highly analogous to cell block cytology sample processing so that the findings from our study would be relevant to oncopanel sequencing of clinical FFPE samples. We identified multiple previously unrecognized and avoidable sources of variation that, if addressed by appropriate QC measures, should enable more reliable use of FFPE samples.

We leveraged results from the SEQC2 Oncopanel Sequencing working group that extensively sequenced this cell line in a companion study^1^ to create variant reference sets. However, the variant reference set for AZ650 was generated from FFPE samples that passed stringent QC filters because AZ650 was not included in the companion study of multiple oncopanels with fresh gDNA samples. This might have led to an underestimation of FP rates for samples sequenced by AZ650 because few variant calls due to FFPE damage may have been included as reference variants. This did not alter our overall conclusion based on the analysis results of FPR because FPR was drastically different for any panel in the pairwise comparisons of genomic regions (**Fig. 4A**), QC status, and sample groups (**Fig. 5A**).

Given that a QC related issue in any single step of FFPE processing, sample preparation, library preparation, sequencing, and/or bioinformatics can increase false positive/negative rates, we sought to survey QC checks throughout the entire process. For the first step (pre-analytical QC), cell count and DNA input were key drivers of the FPR. For the second step (Oncopanel NGS QC), there was a strong and significant correlation between minimum deduplicated sequencing depth (DSD) and FPR. In support of this finding, experiments were far more likely to fail if library complexity was below 0.25. After removing contaminated samples, a minimum median DSD of 600X effectively controlled FPR and separated high quality FFPE samples from failed samples for the three capture-based panels. For each hybrid-capture panel, we observed some moderate correlation between median DSD and DNA input amount for library preparation. However, there was no correlation between cell count and DNA extraction yield for any panel after excluding those surface samples with very low cell count. The loss of correlation may have been due to variations in cell count estimation and DNA extraction efficiency. For the third step (post-variant call QC), G:C to T:A transversions may be employed to pinpoint outliers associated with oxidative damage. A per sample VAF histogram also provided an important QC metric. A long tail from the left side of 100% VAF was likely an indicator of sample contamination. This sample contamination check is applicable to impure tumor samples that may contain DNA from stromal cells. While the exact QC cutoffs employed in this study may not extrapolate perfectly to different panels, tissue samples, and FFPE processes, we expect these QC metrics to remain relevant across oncopanels.

Not all variants fall into equally easy-to-call genomic regions, thus we queried the impact of variant location on FFPE sample calling. All panels generally showed the lowest FPRs in the CTR. Thus, restriction of variant calls to within the CTR boosts reliability in the context of FFPE samples. Genomic region also showed interactions with other QC metrics. For variants falling outside of the CTR, the impact of alternate read depth cutoff was more pronounced (lower alternate read depth cutoff led to higher FPR). In the case of precious clinical samples from an individual patient, it may not always be possible to employ such rigorous QC metrics because choices are limited. Here, we found that it was possible to reduce the FPR through higher VAF cutoff, variant allele depth cutoff, and restriction of variant calling to the consensus high confidence region. The trade-off was reduced sensitivity.

Interestingly, physical position within the FFPE block showed a significant effect on the amount of measured FFPE damage and technical artifacts. Analogous to this, “edge effect” in histologic tissue section staining is observed regularly by pathologists as a source of error in prognostic and treatment marker interpretation^26^. Its cause is not completely understood but is thought to stem from the surfaces of tissue sections experiencing a combination of drying artifact, oxidative damage, as well as formalin fixation damage. In our study, sections taken from the inner portion of the FFPE block were less likely to fail our QC metrics than samples from the surface portions, which almost always failed QC; perhaps due to the same underlying factors causing “edge effect” seen in traditional histologic markers. This leads to an actionable recommendation that samples from the surface portions of an FFPE block should be avoided if possible when selecting samples for oncopanel sequencing. Follow-up studies should be conducted to confirm and further characterize this newly identified intra-sample source of FFPE quality variation. In addition, samples that underwent longer formalin fixation times were more likely to produce less high quality FFPE sections suitable for genomic sequencing. To the extent possible, shortening formalin fixation time is recommended to enhance the quality of genomic sequencing. Importantly, our study employed cell culture samples rather than surgically cut tissue samples. Intact tissue samples harbor widely varying formalin penetration rates, and this can greatly impact formalin fixation time (e.g. brain [fatty] vs muscle [high water content]). For clinical samples, it is important to make samples as thin as possible to improve uniformity of fixation in the shortest amount of time possible. Our findings may not extend quantitatively to FFPE samples from intact tissue samples, but we anticipate our findings will apply qualitatively. Currently there are no clear guidelines for a specimen formalin fixation window to optimize FFPE samples for NGS testing. A recent publication resulted from the CAP Preanalytics for Precision Medicine Project which recommended that specimen sample thickness be less than 5 mm and total fixation time be between 6 and 24 hours for nonfatty tissues to ensure molecular integrity of cancer specimens^4^. To extend this recommendation, we suggest that the total fixation time be limited to 6 hours and the surface portions be avoided when choosing FFPE sections for oncopanel sequencing.

Taken together, our work advocates for a robust set of QC metrics querying various steps in the process from sample to sequencing to bioinformatics. For the first time, through comprehensive multi-laboratory oncopanel sequencing of 96 samples created under well-controlled FFPE processing, we quantitatively evaluated the effects of formalin fixation duration and within-block position on data quality. We specified the portion of FFPE block that would be suitable for oncopanel sequencing experiments and discovered that the size of such portion was dependent on formalin fixation time. Thus, shorter fixation time is recommended to the extent possible. To ensure reliable results, our results support the application of strict threshold criteria for cell count, DNA input, allele frequency, and restriction of analysis to genomic regions of high confidence. FFPE samples are archived on a routine basis in pathology departments around the world. By identifying specific quality control factors that affect targeted NGS analysis of FFPE samples we hope to increase their value in research and clinical diagnostics.

## Materials and Methods

### Sample Preparation, Quality Check, and Distribution

The Agilent male lymphoblast cell line was cultured in T75 flasks (Corning Catalog No. 10-126-28) and harvested according to vendor product specifications. The harvested cellular material was combined into a single 15 mL conical tube (Falcon Catalog No. 14-959-53A) and resuspended to a total volume of 1 mL with 10% neutral buffered formalin (NBF) (StatLab Catalog No. 28600). In separate vials, HistoGel specimen processing gel matrix (ThermoFisher Catalog No. HG-4000-012) was heated to 60 °C for 2 hours to liquify and then allowed to cool and equilibrate to 45 °C in a vendor supplied thermal block (ThermoFisher Catalog No. HGSK-2050-1). Replicate square shape cell-block molds were set up (Fisherbrand Catalog No. EDU00553), and into each mold, 500 μLof 45 °C HistoGel was added; Then, 100 μL of NBF-suspended cell line mixture was added. These were immediately and gently stirred to ensure homogeneity of cells within the cooling HistoGel matrix, and then they were allowed to sit and solidify on the bench top for at least 5 minutes. Next, for each mold, a micro-spatula was used to carefully dislodge the formed HistoGel embedded cell mixtures, and these were carefully placed into nylon mesh bags (ThermoFisher Scientific Catalog No. 6774010) to prevent disaggregation during subsequent tissue processing. Each cell mixture block was 2.67 mm thick with a square cross section of 225 mm^2^. These formed HistoGel cell mixtures in nylon bags were placed into individual tissue processing cassettes (ThermoFisher Scientific Catalog No. 1000957), and then were submerged in a plastic pail filled with 10% NBF to simulate pre-tissue-processing time-in-formalin delay before batch tissue processing steps.

The sequence described above was performed at 1-, 2-, 6-, and 24-hour time points prior to batch tissue processing. All cassettes were then placed into a tissue processor for a “routine” tissue processing run at the University of Toledo Medical Center Department of Pathology (Sakura Tissue Tek VIP 5 Tissue Processor). The processor with 14 stations was programmed as follows: 1) 10% NBF for 1 hour, 2) 10% NBF for 1 hour, 3) 70% ethanol for 1 hour, 4) 80% ethanol for 1 hour, 5) 95% ethanol for 45 minutes, 6) 95% ethanol for 45 minutes, 7) 100% ethanol for 45 minutes, 8) 100% ethanol for 45 minutes, 9) xylene for 45 minutes, 10) xylene for 45 minutes, 11) paraffin at 60 °C for 30 minutes, 12) paraffin at 60 °C for 30 minutes, 13) paraffin at 60 °C for 30 minutes, and 14) paraffin at 60 °C for 0 minutes. The processed formalin fixed paraffin infiltrated cell blocks were then embedded in paraffin (Sakura Tissue Tek TEC 5 Tissue Embedding Station) to create formalin-fixed paraffin embedded (FFPE) cell blocks.

Immunohistochemistry of FFPE materials was prepared using Pan Keratin (Ventana Catalog No. 760-2135) and CONFIRM-anti-CD45 (Ventana Catalog No. 760-2505) reagents using a Benchmark Ultra Ventana Automated IHC slide staining system. These two IHC stains were used to ensure purity of cell cultures after tissue processing.

Each FFPE cell block was serially sectioned at 5 μm thickness with a microtome. Smaller groups of 8 sections (“curl samples”) were placed into individual low-binding Eppendorf tubes for intra-block comparison of FFPE sampling variation. For cell count and quality control purposes, one alternating section was taken for routine hematoxylin and eosin (H&E) staining. Alternating sets of one H&E slide and 8 sections of FFPE material (“curl sets”) were sectioned until the block was exhausted. The relative position “number” was recorded for each H&E slide and used to mark the corresponding “curl-set” tube. To estimate the cellularity for each tube containing formalin fixed and paraffin embedded materials, the average cell count was first computed for the two flanking H&E slides and multiplied by four as a human lymphoblast cell (10-20 μm in size) would likely appear in two adjacent sections of 5 μm thickness. Most cell count estimates ranged from approximately 10,000 to 40,000.

Samples with obviously low counts at the top end of each block were excluded prior to the first recording of relative position. Based on the relative positions and cell count estimates, samples were grouped into three categories: surface (top or bottom) samples with cell count estimates below 50% of the average count per sample, the adjacent surface samples with similar cell counts, and the inner samples. Most inner samples showed cell counts above 20,000. Three adjacent samples were assigned to a second surface category at each end of an FFPE cell block. Occasionally one or two more adjacent samples were assigned to the second surface category if their cell counts were much closer to the surface samples than the inner samples. Samples of the first surface category usually showed cell counts below 10,000. Cell counts for all available samples and their categories are provided in **Supplementary File 1**. All samples were coded by a concatenated string of three fields separated by an underscore “_”: up to two digits for formalin fixation time in hours, a letter in uppercase for FFPE block (“G” or “H”), and up to two digits for FFPE block position. For each formalin fixation time, two FFPE blocks were used, and three samples were taken evenly from each block. A set of 24 FFPE curl samples were then distributed to each testing laboratory.

### Cross-panel targeted NGS testing of FFPE samples

Four laboratories participated in this study and each tested one distinct oncopanel with support from the panel provider. These four panels were AstraZeneca 650 genes Oncology Research Panel (AZ650), Burning Rock DX OncoScreen Plus (BRP), Illumina TruSight Tumor 170 (ILM), and Thermo Fisher Oncomine Comprehensive Assay v3 (TFS). Each laboratory extracted and quantified genomic DNA from the FFPE sections. NGS sequencing experiments were conducted following vendor recommended protocols, with QC data collected as well. Sequencing data was then processed by the respective oncopanel vendor recommended bioinformatics pipeline. Detailed information regarding targeted NGS experiment and variant calling method is provided below for each panel. Variant calling results and QC data were collected and submitted to the Working Group for integrated analysis.

### AZ650 panel NGS testing and variant calling

The AZ650 assay is a hybrid capture panel designed by AstraZeneca to perform next-generation sequencing on solid tumors for exploratory evaluation of pan-cancer biomarkers. AZ650 was designed with reference sequences from human genome HG38. DNA Extraction was performed on FFPE tissue using the Omega M6958 Kit performed on the KingFisher Flex instrument. Extracted DNA was quantitated using the Qubit dsDNA High Sensitivity Kit (cat # Q32854). Each sample was quantitated in duplicate 2 plreactions, and the average was calculated as the final DNA concentration (ng/μL).

DNA whole genome libraries were constructed using the Kapa Biosystems HyperPlus kit (cat # 07962428001, Roche) onboard the Beckman Coulter Biomek FxP liquid handling platform with an integrated on-deck Biometra TRobot thermal cycler. A DNA aliquot was normalized for each sample in 10 mM TRIS-HCl buffer. Enzymatic fragmentation was performed to shear DNA prior to adapter ligation. Unique dual-indexed adapters containing a 6 base-pair UMI sequence were ligated to the fragmented DNA. The DNA whole genome libraries were quantitated using the Agilent TapeStation D1000 (cat # 5067-5582, Agilent). Quantitation values and fragment lengths sourced from the TapeStation D1000 were used for quality control prior to hybridization capture reaction.

Hybridization capture was performed to enrich for the regions of the genome that comprise the targeted panel. Prior to hybrid capture, whole genome libraries were multiplexed together in equimolar ratios, and concentrated using a SPRI bead method. The hybridization capture protocol was performed manually using probes produced by IDT and the Roche NimbleGen SeqCap Hybridization and Wash Kit. Hybrid capture libraries were quantitated using both the Agilent TapeStation D1000 ScreenTape and the Kapa Biosystems Library Quantification kit (qPCR).

Sequencing of each hybrid capture pool was performed on either the Illumina HiSeq 4000 or NovaSeq 6000 sequencers. Each pool was normalized to 1 nM and quantitated via TapeStation D1000, then diluted to a final concentration of 200 pM prior to flowcell loading. Sequencing Analysis Viewer (SAV) and the MultiQC tools^27^ were used to review the quality metrics generated from the sequencer. Sequencing data was demultiplexed, passed through a bcl-to-fastq conversion program^28^ (bcl2fastq v2.20.0.422). FASTQ files were analyzed using pipeline software bcbio-nextgen^29^. Reads were aligned to the hg38 reference using bwa mem v0.7.17^30^, and sequencing duplicates for each UMI were collapsed into a single consensus read using fgbio^31^ v1.0.0. Variant calling was performed using VarDict v1.7.0^32^, down to a variant allele frequency (VAF) of 1% (before filtering and curation) and variant effects annotated by snpEff v4.3.1t^33^. All software was run using best practice parameters established within the bcbio workflow or in-house. Mapped UMI consensus reads (in BAM files) and variant calling results (in VCF files) were then provided to the working group for further analysis. The following variant filters were recommended by the panel provider to minimize false positive calls: 1) a total depth threshold of 100; 2) at least four forward, four reverse, and ten total support reads for the alternative allele; 3) VAF threshold of 2%; and 4) mean position in support reads (pMEAN)^32^ greater than 15.

### BRP panel NGS testing and variant calling

The DNA from 24 individual formalin fixed cell pellets in paraffin scrolls were extracted using the AllPrep DNA/RNA FFPE kit (Qiagen) following the manufacturer’s genomic DNA purification protocol. The extraction process involved deparaffinization, protease digestion, DNA-containing pellet separation, second protease digestion, de-crosslinking, column binding, washing, and elution. After purification, the DNA concentration was quantified using a Qubit Fluorometer with dsDNA HS assay kit (Life Technologies, Carlsbad, CA). The library prep and enrichment process were performed using a Burning Rock HS library preparation kit. The procedure was described previously^1^. In brief, DNA shearing was performed on each FFPE DNA samples using a Covaris M220 for 240s, with Peak Incident Power=50W, Duty Factor: 20%, Cycle Per Burst: 200, at 2-8 °C, followed by end repair, adaptor ligation, and PCR enrichment. Approximately 750 ng of purified pre-enrichment library was hybridized to the OncoScreenPlus panel and further enriched following manufacturer instructions. The OncoScreenPlus panel is approximately 1.7M bp in size and covers 520 human cancer related genes. Final DNA libraries were quantified using a Qubit Fluorometer with dsDNA HS assay kit (Life Technologies, Carlsbad, CA). Agilent 4200 TapeStation D1000 Screen Tape was then performed to assess the quality and size distribution of the library. The libraries were sequenced on a NovaSeq 6000 instrument (Illumina, Inc., San Diego, CA) with 2 × 150 bp pair-end reads with unique dual index.

After demultiplexing using bcl2fastq v2.20^28^ (Illumina), sequence data were filtered using the Trimmomatic0.36^34^ with parameters “TRAILING:20 SLIDINGWINDOW:30:25 MINLEN:50”. Sequence data in fastq format were mapped to the human genome (hg19) using BWA aligner 0.7.10^30^. Local alignment optimization, variant calling and annotation were performed using GATK v3.2.2^35^ with parameters “--interval_padding 100 - known 1000G_phase1.indels.b37.vcf - known Mills_and_1000G_gold_standard.indels.b37.vcf” and VarScan v2.4.3^36^ with parameters “-min-coverage 50 --min-var-freq 0.005 --min-reads2 5 --output-vcf 1 --strand-filter 0 --variants 1 --p-value 0.2”. For SNV and small indels, variants were further filtered using an in-house variant filter pipeline. For each valid variant, the covered raw depth was required to be greater or equal than 50 (DP>=50), and at least 5 mutation supporting count (AD>=5); minor allele frequency was required to be greater than 0.01 (AF>=0.01). In order to further filter out false positives, only variants with at least 6 unique fragments support or 2 unique paired fragment support, i.e., within overlapping region between read pairs, were kept. After filtering, remaining valid variants were annotated with ANNOVAR 20160201^37^ and SnpEff v3.6^33^.

### ILM panel NGS testing and variant calling

FFPE curls were de-paraffinized with xylene followed by an ethanol wash. Briefly, 1 mL xylene was added to each tube with the FFPE sample, vortexed vigorously for 10 seconds, and centrifuged at 20,000 x *g* for 2 minutes. The supernatant was carefully removed. Then 1 mL 96-100% ethanol was added to the pellet, mixed by vortexing, and centrifuged at 20,000 x *g* for 2 minutes. Ethanol was removed and the pellet was air dried. DNA extraction from the pellet was performed using the QIAGEN Allprep DNA/RNA FFPE kit (QIAGEN) on a QIAcube. DNA concentrations were determined by fluorometric quantitation using a Qubit 2.0 Fluorimeter with a Qubit DNA dsDNA HS Assay Kit (Thermo Fisher Scientific).

Library preparation was carried out using the TruSight Tumor 170 Assay (Illumina) following manufacturer’s instructions, except with a lower amount of DNA. Briefly, 30 ng DNA from each sample, except 24G15, 24H10, and 24H15 with 25, 21, and 8 ng DNA respectively, were fragmented on a Covaris Ultrasonicator (Covaris) using the following setting: peak incident power 50 watts, duty factor 30%, cycles per burst 1000, treatment time 270 seconds, and temperature 20 °C. The fragmented DNA was processed through end repair, A-tailing, adapter ligation, and index PCR, enriched by the hybridization-capture method, amplified by final PCR, normalized by bead based normalization, and pooled for sequencing. Twenty-four samples were batched for each library prep, and libraries from every 8 samples were pooled and sequenced on the Illumina NextSeq 550 instrument using the NextSeq High Output Kit. BaseSpace Sequence Hub was used to set up the sequencing run, perform the initial quality control, and generate fastq files for each sample.

Variant calling was performed in BaseSpace Sequence Hub. Briefly, high level sequencing run metrics were evaluated to generate a Run QC Metrics report. Next reads were converted into the fastq format using bcl2fastq^28^, adapters were trimmed and then reads were aligned to the human genome version hg19 using the iSAAC aligner^38^. Indel realignment was performed and then candidate variants were identified using the Pisces variant caller^39^, with a fixed lower limit cutoff for variant allele fraction of at least 2.6%. Variant calls were further compared against a baseline of normal samples to remove systematic false positives.

### TFS panel NGS testing and variant calling

Genomic DNA was extracted from 24 samples of sectioned material using the ALLPrep DNA/RNA FFPE kit (cat #80234). Samples were eluted in 15 μLfrom which 1 μL of material was used for quantification.

Extracted material was prepared for quantification with a Qubit™ dsDNA HS (High Sensitivity) Assay Kit using 1 μL of sample material in 200 μL of Qubit solution. Concentration readings in (ng/μL) from the Qubit Fluorometer were used to calculate 20 ng DNA input in a maximum volume of 7.5 μL into the library preparation.

Libraries were generated using the Ion AmpliSeq Oncomine Comprehensive panel versions 3.0 from Thermofisher Scientific as described^40^ and 17 amplification cycles were performed as suggested for FFPE samples. Final library quantification was performed using real-time PCR (QuantStudio) and the values were given by the instrument in pM. Fifteen out of 24 libraries passed the QC threshold of 50 pM and were successfully sequenced. Five out of 24 libraries did not pass the QC threshold of 50 pM (libraries < 50pM). These samples were cleaned using 1.8 X Ampure beads, eluted in 15 μL then Qubit quantified using 1 μL from the cleaned material. The input DNA amount to be used into library preparation was determined using the samples’ post clean-up Qubit values multiplied by 7.5, which is the maximum volume of input material for library preparation. The newly constructed libraries passed the QC threshold of 50 pM and were successfully sequenced.

The remaining 4 out of 24 libraries did not pass the QC threshold of 50 pM (libraries < 50pM). These samples were cleaned using 1.8 X Ampure beads, eluted in 15 μL then Qubit quantified using 1 μL from the cleaned material. Qubit values were too low to be detected, so 7.5 μL was used as input for library preparation. The newly constructed libraries did not pass the QC threshold of 50 pM and therefore were not sequenced (libraries < 50pM).

All barcoded samples were sequenced on the Ion S5 XL System. Analyses of sequencing raw data were performed with the Ion Torrent Suite (5.10.0). After base calling, reads were aligned with the TMAP module and variant calling was performed with Torrent Variant Caller (TVC), a variant calling module optimized for Ion Torrent data. The default thresholds used for SNVs and indels was 2.5%. Finally, a series of post-calling filters were applied to variant calls to eliminate potential artifacts by fitting the statistical model of flow signals for the observed reference and non-reference alleles. For more details, please see the description of its bioinformatics pipeline in the cross-laboratory oncopanel study^1^ where this panel was used to test a set of reference samples including Agilent male DNA control.

### Identification and Analysis of false positive calls

#### Consensus high confidence targeted region (CTR)

In our collective consortium effort to establish a verified genomic reference material^18^ suitable for assessing the performance of oncology panels in detecting small variants of low allele frequency, a consensus high confidence region for targeted NGS was adopted and later tested in a cross-laboratory oncopanel study^1^ of eight oncology panels where the CTR was shown to reduce the rate of false positive variant calls. The CTR resulted from intersecting exonic coding regions, the NIST high confidence region, and the targeted regions of four whole-exome sequencing panels. Low complexity regions were then removed from the CTR.

#### VCF files cleaning procedures

VCF files provided by each panel vendor were cleaned through a series of procedures. First, each original VCF file was converted into standardized VCF format. INDELs were then normalized with left alignment and trimming using GATK^35^. Complex variants were decomposed with RTG “vcfdecompose”. After that, only variants within the panel regions (excluding the low complexity regions) were kept. Specifically, for VCF files from the TFS panel, we also removed the blacklist variants provided by Thermo Fisher Scientific. To further remove the less confident variants, VAF thresholds recommended by the panel providers were applied for each panel: 2% for AZ650, 1% for BRP, 2.6% for ILM, and 2.5% for TFS. All downstream analyses were applied to the cleaned VCF files.

#### Generation of known variant set for each panel

The known variant set was generated from the fresh DNA samples of each panel. Any variants called in over 75% of all samples were each considered as a known variant. However, due to only three fresh DNA samples that were sequenced for AZ650 panel, instead of using the fresh DNA samples, we adopted high quality FFPE samples with deep sequencing depth (after excluding two contaminated samples) for generation of the known variant set. For the ILM panel, we excluded 6 fresh DNA samples for their low median depth (less than 850). Finally, 20 FFPE samples from AZ650, 12 fresh DNA samples from BRP, 10 fresh DNA samples from ILM, and 12 fresh DNA samples from TFS were used for the known variant set generation for each panel.

#### The false positive estimation and type classification

After the cleaning procedures, any variants called by a panel that were not in the known variant set for that panel were determined as false positive calls. Two metrics were used in the study. The false positive call count was the number of false positive variants for each library, and the false positive rate was then estimated as the ratio of false positive calls out of every million positions defined by each panel. Besides the false positive estimation from the originally cleaned library, false positives could be further reduced by applying more stringent cutoffs. Two types of additional filters were adopted in this study, VAF cutoffs from the panel default VAF threshold to 10% and alternative allele depth cutoffs from 0 to 30. For analysis purposes, besides evaluating the impact of various factors on the FP rate, we also examined the effect per variant type by grouping variant calls into four types: INDELs, G:C to A:T transitions, G:C to T:A transversions, and any other variants.

## Availability of data and materials

DNA-seq data (in FASTQ or BAM format) from four oncopanels will be deposited in the National Center for Biotechnology Information (NCBI) BioProject repository. Variant call results (in VCF format) and dependent files (e.g., panel BED files, genome FASTA files, and others) will be made publicly available through FigShare.

## Acknowledgements

All SEQC2 participants freely donated their time, reagents, and computing resources for the completion and analysis of this project. Part of this work was carried out with the support to Thomas Blomquist through the University of Toledo College of Medicine Academic Affiliation with ProMedica research funds. Leming Shi was supported by the National Key R&D Project of China (2018YFE0201600), the National Natural Science Foundation of China (31720103909), and Shanghai Municipal Science and Technology Major Project (2017SHZDZX01). Donald J. Johann, Jr. acknowledges the support by FDA BAA grant HHSF223201510172C. Nikola Tom was supported by research infrastructure EATRIS-CZ, ID number LM2018133 funded by MEYS CR and MEYS CR project CEITEC 2020 (LQ1601).

The contents of these published materials are solely the responsibility of the administering institution, a participating institution, or individual authors, and they do not reflect the views of any funding body listed above.

## Disclaimer

The views presented in this article do not necessarily reflect those of the U.S. Food and Drug Administration. Any mention of commercial products is for clarification and is not intended as an endorsement.

## List of Supplementary Materials

**Supplementary File 1.xlsx | List of FFPE sample information and extensive quality check data**

**Supplementary File 2.xlsx | List of known variants for four participating oncology panels**

**Supplementary Table 1 | List of detailed information for four participating oncology panels**

**Supplementary Figure 1 | Plot of false positive rate for QC-passed inner FFPE samples showed no observable effect of formalin fixation time.** QC-passed inner FFPE samples were aggregated from all four panels after Z-score normalization of false positive rate within the sample group per panel. Within each panel’s sample group, the mean and standard deviation of the false positive rate were first calculated. Z-scored false positive rate was calculated by dividing its difference from the mean by the standard deviation. This normalization step removed the differences in false positive rate between panels. Samples were then grouped by formalin fixation time (x-axis) and plotted together. Each circle represents a sample with its normalized false positive rate shown on the y-axis. Samples are color coded per panel: red for AZ650, green for BRP, blue for ILM, and black for TFS. There was no observable effect of formalin fixation time on false positive rate.

## References

1. SEQC2 Oncopanel Sequencing Working Group. Cross-oncopanel study reveals high sensitivity and accuracy with overall analytical performance depending on genomic regions. Accept. Genome Biol. GBIO--20-01201.

2. Jennings, L. J. et al. Guidelines for Validation of Next-Generation Sequencing–Based Oncology Panels: A Joint Consensus Recommendation of the Association for Molecular Pathology and College of American Pathologists. J. Mol. Diagn. 19, 341–365 (2017).

3. Agrawal, L., Engel, K. B., Greytak, S. R. & Moore, H. M. Understanding Preanalytical Variables and their Effects on Clinical Biomarkers of Oncology and Immunotherapy. Semin. Cancer Biol. 52, 26–38 (2018).

4. Compton, C. C. et al. Preanalytics and Precision Pathology: Pathology Practices to Ensure Molecular Integrity of Cancer Patient Biospecimens for Precision Medicine. Arch. Pathol. Lab. Med. 143, 1346–1363 (2019).

5. AJCC Cancer Staging Manual, 8th edition. (Springer International Publishing, 2017).

6. Wolff, A. C. et al. Recommendations for Human Epidermal Growth Factor Receptor 2 Testing in Breast Cancer: American Society of Clinical Oncology/College of American Pathologists Clinical Practice Guideline Update. Arch. Pathol. Lab. Med. 138, 241–256 (2014).

7. Hammond, M. E. H. et al. American Society of Clinical Oncology/College of American Pathologists guideline recommendations for immunohistochemical testing of estrogen and progesterone receptors in breast cancer (unabridged version). Arch. Pathol. Lab. Med. 134, e48–72 (2010).

8. Minimally Invasive Surgery Market-Global Industry Analysis, Size, Share, Growth, Forecast 2019. https://www.transparencymarketresearch.com/minimally-invasive-surgery-market.html.

9. Do, H. & Dobrovic, A. Sequence Artifacts in DNA from Formalin-Fixed Tissues: Causes and Strategies for Minimization. Clin. Chem. clinchem.2014.223040 (2014) doi:10.1373/clinchem.2014.223040.

10. Decision Memo for Next Generation Sequencing (NGS) for Medicare Beneficiaries with Advanced Cancer (CAG-00450N). 151.

11. Centers for Medicare & Medicaid. National Coverage Determination (NCD) for Next Generation Sequencing (NGS) (90.2). https://www.cms.gov/medicare-coverage-database/details/ncd-details.aspx?NCDId=372.

12. FoundationOne. FoundationOne CDx Technical Information.

13. Lott, R. et al. Practical Guide to Specimen Handling in Surgical Pathology. 53 (2015).

14. College of American Pathologists. Anatomic Pathology Checklist, College of American Pathologists. (2017).

15. Wong, S. Q. et al. Sequence artefacts in a prospective series of formalin-fixed tumours tested for mutations in hotspot regions by massively parallel sequencing. BMC Med. Genomics 7, 23 (2014).

16. Van Allen, E. M. et al. Whole-exome sequencing and clinical interpretation of formalin-fixed, paraffin-embedded tumor samples to guide precision cancer medicine. Nat. Med. 20, 682–688 (2014).

17. Aziz, N. et al. College of American Pathologists’ Laboratory Standards for Next-Generation Sequencing Clinical Tests. Arch. Pathol. Lab. Med. 139, 481–493 (2015).

18. Jones, W. D. & SEQC2 WG2 group. A Validated Genomic Reference Material for Assessing Performance of Cancer Panels Detecting Small Variants of Low Allele Frequency. Genome Biol. Submiss. ID GBIO--20-01193.

19. Zook, J. M. et al. Integrating human sequence data sets provides a resource of benchmark SNP and indel genotype calls. Nat. Biotechnol. 32, 246–251 (2014).

20. Fang, L. T. & SEQC2 WG1 working group. Establishing reference samples for benchmarking detection of somatic mutations germline and variants with NGS technologies. Nat. Biotechnol. NBT-RS47789 Rev.

21. Van Ooyen, S., Loeffert, D. & Korfhage, C. Overcoming constraints of genomic DNA isolated from paraffin-embedded tissue. Qiagen White Pap. 1–6.

22. Spencer, D. H. et al. Comparison of Clinical Targeted Next-Generation Sequence Data from Formalin-Fixed and Fresh-Frozen Tissue Specimens. J. Mol. Diagn. 15, 623–633 (2013).

23. Sekiguchi, M. & Tsuzuki, T. Oxidative nucleotide damage: consequences and prevention. Oncogene 21, 8895–8904 (2002).

24. Do, H. & Dobrovic, A. Sequence Artifacts in DNA from Formalin-Fixed Tissues: Causes and Strategies for Minimization. Clin. Chem. 61, 64–71 (2015).

25. Prentice, L. M. et al. Formalin fixation increases deamination mutation signature but should not lead to false positive mutations in clinical practice. PLOS ONE 13, e0196434 (2018).

26. Gown, A. M. Current issues in ER and HER2 testing by IHC in breast cancer. Mod. Pathol. 21, S8–S15 (2008).

27. Ewels, P. MultiQC: Summarize results from bioinformatics analysis across many samples into a single report. Bioinformatics 32(19), 3047–8 (2016).

28. Illumina. bcl2fastq2 Conversion Software v2.20. https://support.illumina.com/downloads/bcl2fastq-conversion-software-v2-20.html.

29. Brad Chapman et al. bcbio/bcbio-nextgen: v1.2.4. (Zenodo, 2020). doi:10.5281/zenodo.4041990.

30. Li, H. Aligning sequence reads, clone sequences and assembly contigs with BWA-MEM. ArXiv13033997 Q-Bio (2013).

31. fulcrumgenomics, fulcrumgenomics, & fulcrumgenomics. fgbio. (Fulcrum Genomics, 2020).

32. Lai, Z. et al. VarDict: a novel and versatile variant caller for next-generation sequencing in cancer research. Nucleic Acids Res. 44, e108 (2016).

33. Cingolani, P. et al. A program for annotating and predicting the effects of single nucleotide polymorphisms, SnpEff. Fly (Austin) 6, 80–92 (2012).

34. Bolger, A. M., Lohse, M. & Usadel, B. Trimmomatic: a flexible trimmer for Illumina sequence data. Bioinformatics 30, 2114–2120 (2014).

35. McKenna, A. et al. The Genome Analysis Toolkit: A MapReduce framework for analyzing next-generation DNA sequencing data. Genome Res. 20, 1297–1303 (2010).

36. Koboldt, D. C. et al. VarScan 2: Somatic mutation and copy number alteration discovery in cancer by exome sequencing. Genome Res. 22, 568–576 (2012).

37. Wang, K., Li, M. & Hakonarson, H. ANNOVAR: functional annotation of genetic variants from high-throughput sequencing data. Nucleic Acids Res. 38, e164–e164 (2010).

38. Raczy, C. et al. Isaac: ultra-fast whole-genome secondary analysis on Illumina sequencing platforms. Bioinformatics 29, 2041–2043 (2013).

39. Dunn, T. et al. Pisces: an accurate and versatile variant caller for somatic and germline next-generation sequencing data. Bioinformatics 35, 1579–1581 (2019).

40. Pain, M. et al. Treatment-associated TP53 DNA-binding domain missense mutations in the pathogenesis of secondary gliosarcoma. Oncotarget 9, 2603–2621 (2017).

