## Supplementary figures and images for "Deep oncopanel sequencing reveals fixation time- and within block position-dependent quality degradation in FFPE processed samples"

### Supp figure 1

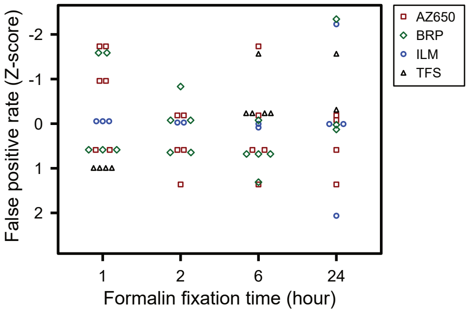
