## Supplementary material for "Deep oncopanel sequencing reveals fixation time- and within block position-dependent quality degradation in FFPE processed samples": Supp table 1

### Table S1 | List of detailed information for four participating oncology panels

| Panel code | Panel name | Genome version | Gene count | Design Region (Kbp) | Reporting Region (Kbp) |
|---|---|---|---|---|---|
| AZ650 | AstraZeneca 650 genes Oncology Research Panel | hg38 | 650 | 1,808 | 1,808 |
| BRP | Burning Rock DX OncoScreen Plus | hg19 | 523 | 1,631 | 1,072 |
| ILM | Illumina TruSight Tumor 170 | hg19 | 154 | 527 | 527 |
| TFS | Thermo Fisher Oncomine Comprehensive Assay v3 | hg19 | 146 | 349 | 289 |

*continue…*

| Panel code | Reporting Region within the CTR* (Kbp) | Reporting Region outside the CTR (Kbp) | Fragmentation approach |
|---|---|---|---|
| AZ650 | 1116 | 692 | Enzymatic fragmentation |
| BRP | 823 | 250 | Covaris M220/Focused-Ultrasonicators with AFA Technology |
| ILM | 349 | 178 | Covaris / Focused-Ultrasonicators |
| TFS | 174 | 115 | No fragmentation as PCR based target amplification |

*continue…*

| Panel code | Enrichment | Sequencing platform | Read length | UMI | Avg. read count |
|---|---|---|---|---|---|
| AZ650 | capture based | HiSeq 4000 or NovaSeq 6000 | 2 x 150bp | Yes | 40.8M (consensus reads) |
| BRP | capture based | NovaSeq 6000 | 2 x 150bp | No | 74.3M |
| ILM | capture based | NextSeq 550 | 2 x 101bp | No | 80.7M |
| TFS | amplicon based | IonTorrent S5 XL | 113 bp (average) | No | 15.4M (mapped) |

*continue…*

| Panel code | Read mapping tool | Variant caller | VAF threshold |
|---|---|---|---|
| AZ650 | bwa mem v0.7.17 | fabio v1.0.0 and VarDict v1.7.0 | 2% |
| BRP | BWA aligner 0.7.10 | VarScan v2.4.3 | 1% |
| ILM | iSAAC aligner | Pisces variant caller | 2.6% |
| TFS | TMAP | Torrent Variant Caller (TVC) | 2.5% |

* CTR is the consensus high confidence targeted region (see Methods for details).
